# Binding of His-tagged fluorophores to lipid bilayers of giant vesicles†

**DOI:** 10.1101/2022.02.01.478643

**Authors:** Shreya Pramanik, Jan Steinkühler, Rumiana Dimova, Joachim Spatz, Reinhard Lipowsky

**Affiliations:** Max Planck Institute of Colloids and Interfaces, 14424 Potsdam,Germany; Department of Biomedical Engineering, Northwestern University, Evanston, IL, USA; Max Planck Institute for Medical Research, 69120 Heidelberg, Germany

## Abstract

His-tagged molecules can be attached to lipid bilayers via certain anchor lipids, a method that has been widely used for the bio-functionalization of membranes and vesicles. To observe the membrane-bound molecules, it is useful to consider His-tagged molecules that are fluorescent as well. Here, we study two such molecules, green fluorescence protein (GFP) and green-fluorescent fluorescein isothiocyanate (FITC), both of which are tagged with a chain of six histidines (6H) that bind to the anchor lipids within the bilayers. The His-tag 6H is much smaller than the GFP molecule but somewhat larger than the FITC dye. The lipid bilayers form giant unilamellar vesicles (GUVs), the behavior of which can be directly observed in the optical microscope. We apply and compare three well-established preparation methods for GUVs: electroformation on platinum wire, polyvinyl alcohol (PVA) hydrogel swelling, and electroformation on indium tin oxide (ITO) glass. Microfluidics is used to expose the GUVs to a constant fluorophore concentration in the exterior solution. The brightness of membrane-bound 6H-GFP exceeds the brightness of membrane-bound 6H-FITC, in contrast to the quantum yields of the two fluorophores in solution. In fact, 6H-FITC is observed to be strongly quenched by the anchor lipids which bind the fluorophores via Ni^2+^ ions. For both 6H-GFP and 6H-FITC, the membrane fluorescence is measured as a function of the fluorophores’ molar concentration. The theoretical analysis of these data leads to the equilibrium dissociation constants *K*_*d*_ = 37.5 nM for 6H-GFP and *K*_*d*_ = 18.5 nM for 6H-FITC. We also observe a strong pH-dependence of the membrane fluorescence.

## 1 Introduction

Lipid bilayers, which represent a universal building block for all biomembranes and, thus, for the architecture of the living cell, provide a versatile biomimetic module to understand the mechanical properties of cells and organelles as well as their responses to external stimuli. In aqueous solutions, lipid bilayers form closed vesicles which have a wide range of applications in the field of synthetic biology ^1–5^, intracellular cargo transport and trafficking ^6–9^ and the systematic study of biomembrane properties ^10–13^. From an experimental point of view, giant unilamellar vesicles (GUVs) are particularly useful because they are micrometer-sized, comparable to the size of eukaryotic cells. In addition, GUVs can be directly imaged by optical microscopy and their mechanical responses can be probed by optical tweezers and micropipettes ^14^.

Two broad categories of membrane proteins can be distinguished. First, integral membrane proteins have one or several transbilayer domains, which span the hydrophobic core of the bilayer membranes. Second, peripheral membrane proteins, which take part in cell signaling and membrane trafficking, can attach to one leaflet of the bilayer membranes by binding to a specific lipid or to a small cluster of several lipids ^15^. Likewise, to achieve biofunctionalisation of GUVs, integral membrane proteins are inserted into the membranes using detergents ^16^ or proteoliposomes ^17,18^ whereas peripheral membrane proteins are attached via certain anchor lipids. Here, we will consider a specific lipid anchor as provided by DGS-NTA(Ni) or NTA lipid for short, ^*^ which binds to poly-histidine tagged molecules via coordinate bonds. Two such molecules will be considered, His-tagged GFP and His-tagged FITC, both of which are green fluorescent, see Figure 1.

**Fig. 1:**
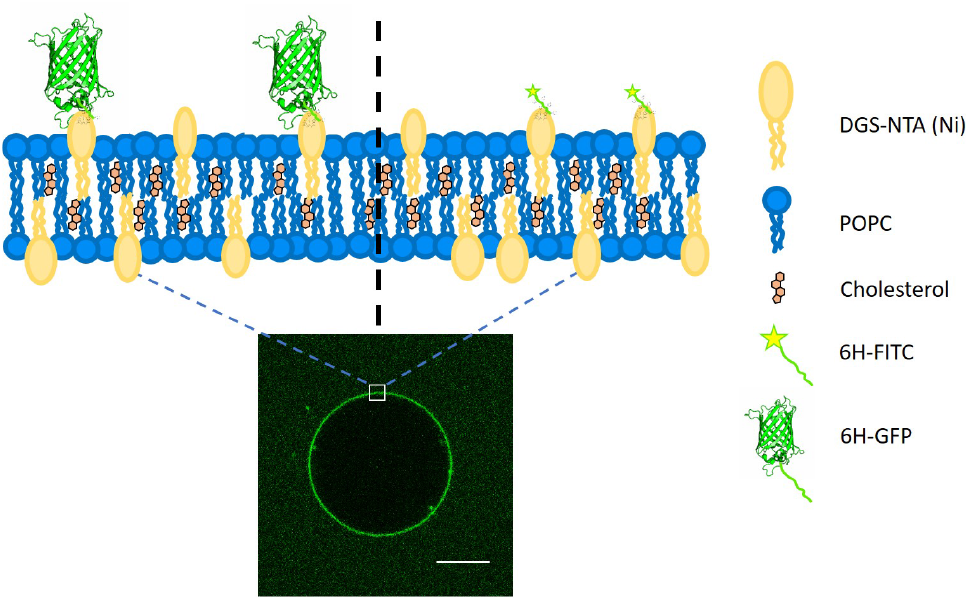
Cartoon depicting the lipid bilayer composed of POPC, cholesterol, and DGS-NTA(Ni) or NTA lipid, which acts as an anchor for a poly-histidine chain. Two types of molecules tagged with a chain of six histidines will be studied, 6H-GFP and the 6H-FITC, both of which are green fluorescent. The barrel height of 6H-GFP is comparable to the membrane thickness (left) whereas the linear extension of 6H-FITC is governed by the length of the histidine chain (right). The optical image displays a GUV with membrane-bound 6H-GFP. Scale bar: 10 *μ*m.

Poly-histidine tags are generally attached to one of the terminals of the proteins for their purification ^19^. Hence, this specific interaction can be harnessed to attach proteins to lipid bilayers ^3,6,20–24^. The NTA(Ni) in the lipid head group forms an octahedral coordinate complex with poly-histidine chains. Four vertices of the octahedron are occupied by the NTA group, leaving two vertices for the binding of two imidazole side chains, see Figure 2. The nitrogen from the imidazole side chain of histidine donates electrons for the coordinate bond. The optimal length of the histidine chain to bind to NTA(Ni) lipids consists of six residues, corresponding to the maximum of the equilibrium association constant as a function of chain length ^20^.

**Fig. 2:**
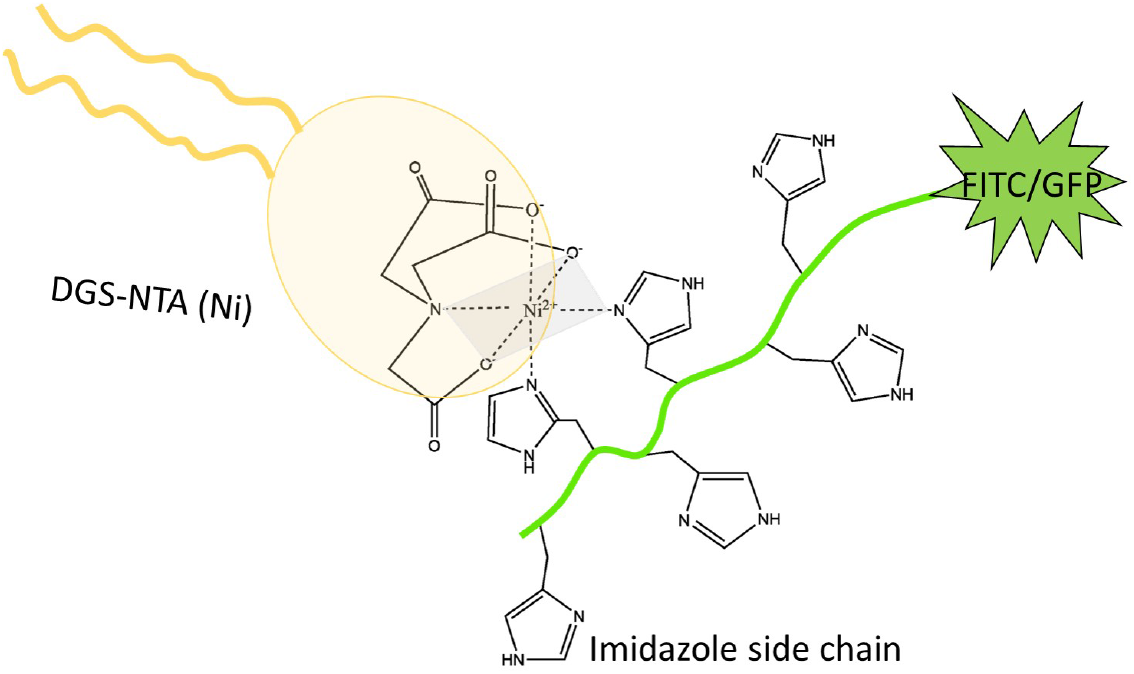
Both 6H-FITC and 6H-GFP consist of a green fluorescent dye attached to a chain of six histidines with six side chains, corresponding to six imidazole groups. Two of the latter groups interact wth the head group of the DGS-NTA(Ni) lipid forming an octahedral complex. Four vertices of this octahedron (grey) are occupied by three oxygens and one nitrogen from the head group, two vertices by two nitrogens from two imidazole side chains. ^20^

The binding between His-tagged proteins and NTA anchor lipids has been frequently used in previous studies. Examples include protein crowding on vesicles ^25^, lipid-coated substrates for drug delivery systems ^26^, artificial cell adhesion ^27^, and high spontaneous curvature generated by the dilute concentration of surface proteins ^28^. In general, the Ni^2+^ ion can be replaced by a Co^3+^ ion in the NTA group. Poly-histidine chains form a stronger and less labile coordinate bond with NTA-Co^3+^ compared to the corresponding bond with NTA-Ni^2+. 29–31^

GUVs can be prepared using a variety of well-established protocols such as swelling lipid layers on a substrate by hydration, electroformation ^32^, phase transfer across liquid-liquid interfaces ^33,34^, and droplet stabilized compartments ^22^; for a recent review, see Ref 14. All of these techniques have their advantages and disadvantages. The swelling methods involve the spreading of a lipid mixture on a substrate followed by the evaporation of the (organic) solvent. As a result, one obtains a stack of lipid bilayers which can then be swollen by hydration. Spontaneous swelling is achieved by spreading the lipid on a non-reactive surface and gently hydrating it for an appropriate period of time. ^35^ This gentle hydration method is limited to the use of specific lipids and solution conditions ^36^.

The influx of the hydrating solution can be enhanced by hydrogel-assisted swelling ^36,37^. The lipid film is deposited on the surface of the dried hydrogel. The hydrogel absorbs the hydrating solution and GUVs form rapidly at the hydrogel-aqueous solution interface. The hydrogels typically used for this purpose are agarose ^37,38^, polyvinyl alcohol (PVA) ^28,36,39^, and polyacrylamide gels ^40^. Sometimes the hydrogel becomes incorporated into the lumen of the GUV or even within the lipid bilayer, which represents a drawback of this method because it affects the mechanical bilayer properties ^38,40^. To avoid these unwanted side effects, electroformation can be used to control the hydration of the lipid films by external alternating currents (ACs) ^32^. The substrate for spreading the lipid film must be electrically conductive. Commonly used substrates include platinum wires ^32,41^, ITO coated glasses ^42–44^, stainless steel wires ^45,46^, copper electrodes ^47^, and carbon fiber microelectrodes ^48^.

One implicit assumption that is often made in the preparation of GUVs is that these vesicles have the same composition as the lipid mixture that was initially used to grow them. This assumption should be checked, especially when the GUVs are prepared from a lipid mixture with a certain fraction of DGS-NTA(Ni). Furthermore, the binding of the poly-histidine with the NTA(Ni) lipid is also sensitive to the solution conditions. Here, we will analyze and compare different preparation techniques of GUVs in different pH conditions. As shown in Figures 1 and 2, we will use two different fluorescent probes, 6H-GFP and 6H-FITC, where 6H stands for a linear chain of six histidines, and study the binding of these fluorophores to NTA lipids embedded in GUV membranes.

The molecular weight of the two fluorophores is quite different: GFP has a molecular weight of 27 kD whereas FITC has a much smaller molecular weight of only 389 D. Therefore, the two fluorophores are also quite different in size. As shown below, one important consequence of this size difference is that the fluorescence of FITC is strongly quenched by the NTA lipids, in contrast to the fluorescence of GFP.

## 2 Results and Discussion

### 2.1 Brightness of GUV membranes

Three different preparation methods are used to form the GUVs: electroformation on platinum wire, PVA hydrogel swelling, and electroformation on ITO glass. In all cases, we start from the same initial lipid mixture with 3 mol% of DGS-NTA(Ni) to grow the GUVs, which are then exposed to a certain concentration of the His-tagged fluorophores. After 10 mins, the GUVs are directly observed with a confocal microscope, see the images in Figure 3.

**Fig. 3:**
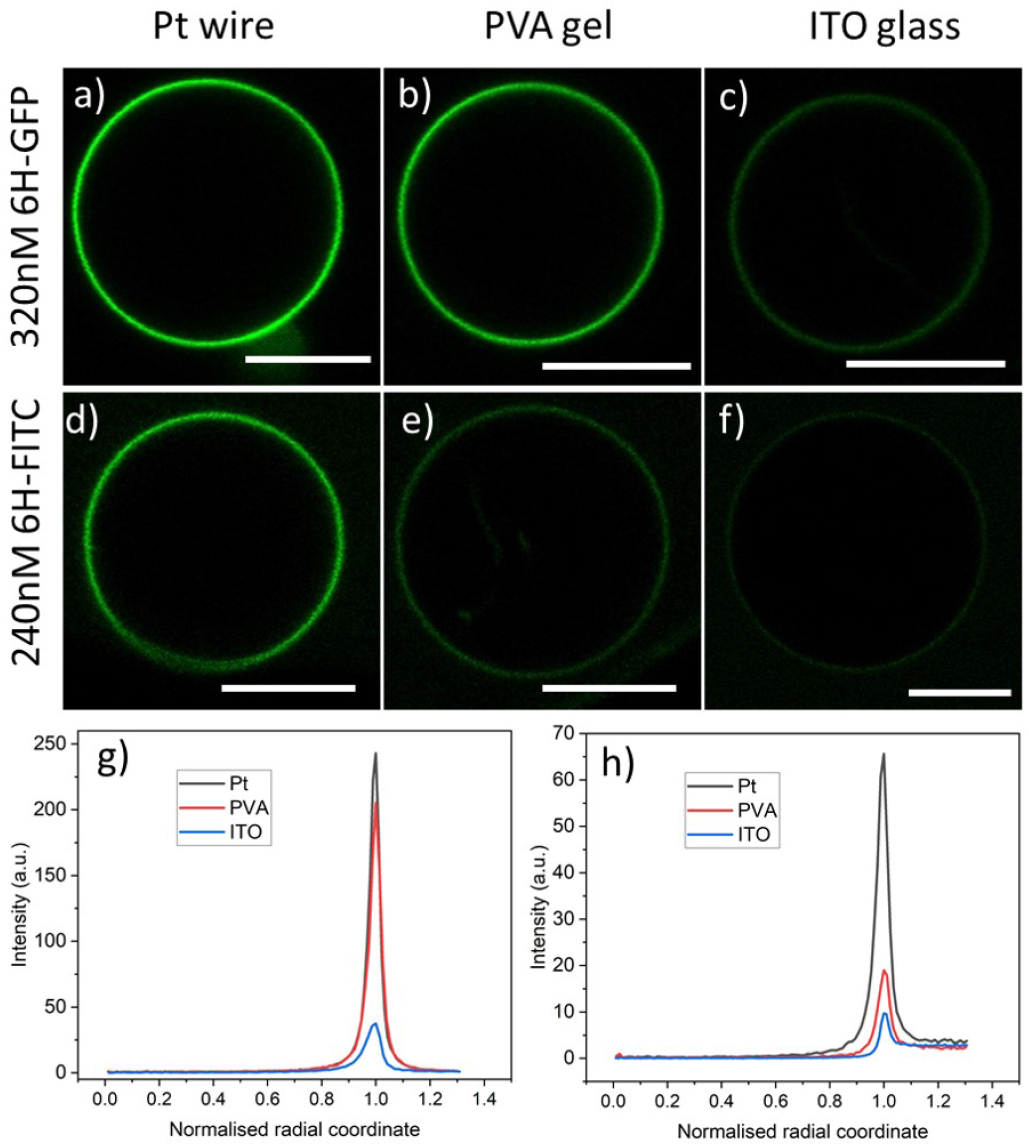
Brightness of GUV membranes doped with DGS-NTA(Ni) for three preparation methods corresponding to the three columns: (a-c) GUV membranes exposed to an exterior solution with 320 nM of 6H-GFP. The Pt wire method leads to a strongly fluorescent membrane whereas this fluorescence is not visible for the PVA gel and ITO glass methods; and (d-f) GUV membranes exposed to an exterior solution with 240 nM of 6H-FITC. The membrane fluorescence is visible for both the Pt wire and the PVA gel methods but invisible for the ITO glass method. The lipid composition used for the GUVs is POPC:Chol (8:2) and 3 mol% DGS-NTA(Ni). In all cases, the GUV encloses a 50 mM sucrose solution and is exposed to an exterior solution of 22.5 mM NaCl and 5 mM sucrose at pH 7.45 with either 6H-GFP or 6H-FITC. All scale bars: 10 *μ*m; and (g,h) Line profiles of fluorescence excess intensity versus normalized radial coordinate; for more details, see Figure 4. The excess intensity profiles in (g) and (h) correspond to the images in (a-c) and (d-f), respectively.

In order to expose the GUVs to a constant concentration of the His-tagged molecules in the exterior solution, we control this concentration by microfluidics and a specific design of the microfluidic chip: We trap individual GUVs in the short side channels of the chip and use its main channel for the fast exchange of the exterior solution. Further details about the microfluidic approach are described in the *Material and Method* section and Figure 10.

Two types of His-tagged molecules are studied: 6H-GFP, corresponding to a chain of six histidines attached to the N terminal of GFP 28, and 6H-FITC consisting of FITC covalently bound to the same histidine chain. The histidine chain binds to the NTA(Ni) head group of the DGS-NTA(Ni) lipid in the vesicle via a coordinate bond (Figure 2).

As shown in Figure 3, the observed brightness of the GUV membranes depends both on the His-tagged fluorophore and on the preparation method. For both fluorophores, the GUVs are observed to have the largest brightness when prepared by platinum wire electroformation. As will become clear further below in Figures 6 and 7, both molar concentrations chosen in Figure 3 – 320 nM for 6H-GFP and 240 nM for 6H-FITC – belong to the saturation regime for the platinum wire method, in which almost all NTA lipids are occupied by His-tagged molecules.

For a spherical GUV, the membrane fluorescence can be determined quantitatively by measuring the fluorescence intensity profile as a function of the radial coordinate *r*, which represents the distance from the center of the vesicle. One example for such a profile is shown in Figure 4a together with the baseline intensity that interpolates smoothly between the background intensities of the interior and exterior solutions. The excess intensity of the fluorescence is equal to the difference between the total fluorescence and baseline intensities, see Figure 4b, and the membrane fluorescence is obtained by integrating this excess intensity over the radial coordinate *r*. The excess intensity profiles corresponding to the GUVs in Figure 3a-f are displayed in Figure 3g,h.

**Fig. 4:**
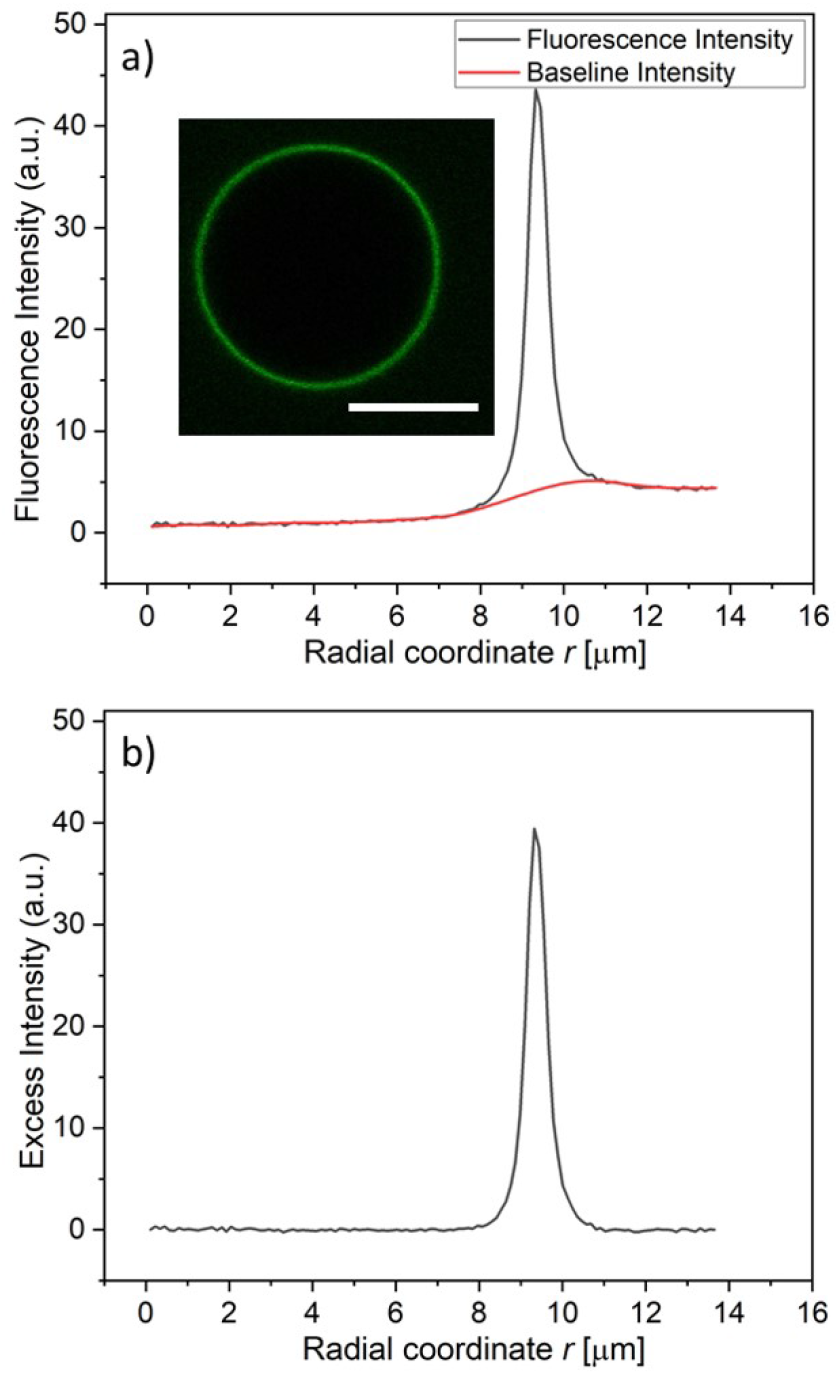
Measurement of membrane fluorescence from His-tagged fluorophores bound to a GUV membrane: (a) Fuorescence intensity (black) and baseline intensity (red) as a function of the radial coordinate *r*, which measures the distance from the vesicle center. The fluorescence intensity was obtained via the image processing software ImageJ, the baseline intensity via OriginLab; and (b) Excess intensity obtained by substracting the baseline intensity from the fluorescence intensity in (a). The membrane fluorescence is obtained by integrating the excess intensity over the radial coordinate *r*. The intensities were obtained for the vesicle displayed in the inset of (a), corresponding to 3 mol% DGS-NTA(Ni), 120 nM 6H-FITC, and pH 7.45. Scale bar: 10 *μ*m.

The GUV images and line profiles show that the brightness of membrane-bound 6H-FITC is reduced compared to the one of membrane-bound 6H-GFP. In contrast, in the absence of lipids, 6H-FITC has a quantum yield that is about 1.2 larger than the quantum yield of 6H-GFP, as obtained from independent measurements, see *Material and Method* section. This different behavior indicates that the fluorescence of 6H-FITC is quenched when this molecule is bound to the membrane or, more precisely, to an NTA anchor lipid.

### 2.2 Fluorescence quenching by NTA lipids

In order to elucidate the quenching effect of the NTA lipids, we added, for each fluorophore, an increasing concentration of NTA lipids to a 120 nM solution of the fluorophore. The resulting fluorescence intensities are displayed in Figure 5. As shown in Figure 5a, the fluorescence of 6H-GFP is hardly affected by large concentrations of NTA lipids, whereas the fluorescence of 6H-FITC is strongly reduced by small concentration of these lipids, see Figure 5b.

**Fig. 5:**
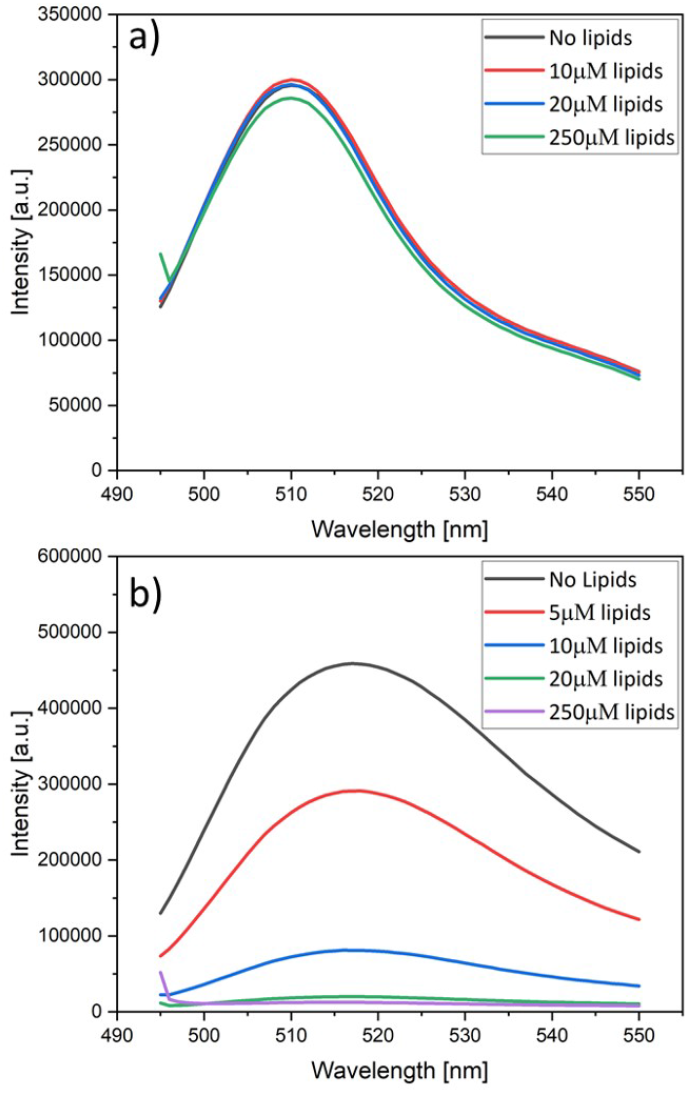
Fluorescence intensity versus wavelength as measured for a 120 nM solution of (a) 6H-GFP and (b) 6H-FITC. Both solutions are exposed to an increasing concentration of lipids in the form of liposomes containing 30 mol% NTA lipids, see color codes in the insets. The fluorescence intensity of 6H-GFP in (a) is hardly affected by the lipids whereas the fluorescence intensity of 6H-FITC in (b) is strongly reduced.

The data in Figure 5a,b were obtained by adding liposomes with the lipid composition POPC:Chol:NTA as given by 6:1:3. Similar quenching effects were observed when the fluorophores was exposed to nickel sulphate. On the other hand, no fluorescence quenching of 6H-FITC was observed when the liposomes contained only POPC and cholesterol but no NTA lipids, as demonstrated in Figure S1.

Therefore, the fluorescence quenching of 6H-FITC is caused by NTA lipids which bind the fluorophore via their Ni^2+^ ion, see Figure 2. Fluorescence quenching by Ni^2+^ ions has been previously reported for a variety of fluorophores. ^49–51^ For membrane-bound 6H-GFP, such a quenching effect does not occur because the chromophore of 6H-GFP is located in the middle of the protein’s *β*-barrel ^52^ and well-separated from the Ni^2+^ ion of the NTA lipid. In contrast, the fluorescein group of membrane-bound 6H-FITC is much closer to this Ni^2+^ ion, see Figure 2, which explains the strong quenching effect in Figure 5b.

### 2.3 Fluorescence of membrane-bound 6H-GFP

Using the Pt wire method, we measured the membrane fluorescence *I*_flu_ of 6H-GFP as a function of its molar concentration *X* for the range 0 ≤ *X* ≤ 320 nM. The corresponding normalised intensity *I*_flu_*/I*_sat_ is plotted in Figure 6a with *I*_sat_ = 320.4 a.u. where a.u. stands for arbitrary (or procedure-defined) units. These data are well fitted by the intensity-concentration relationship as given by eqn (6) in the *Material and Method* section. From this fit, we obtain the equilibrium dissociation constant *K*_*d*_ = (37.5 ± 7.5) nM for 6H-GFP.

**Fig. 6:**
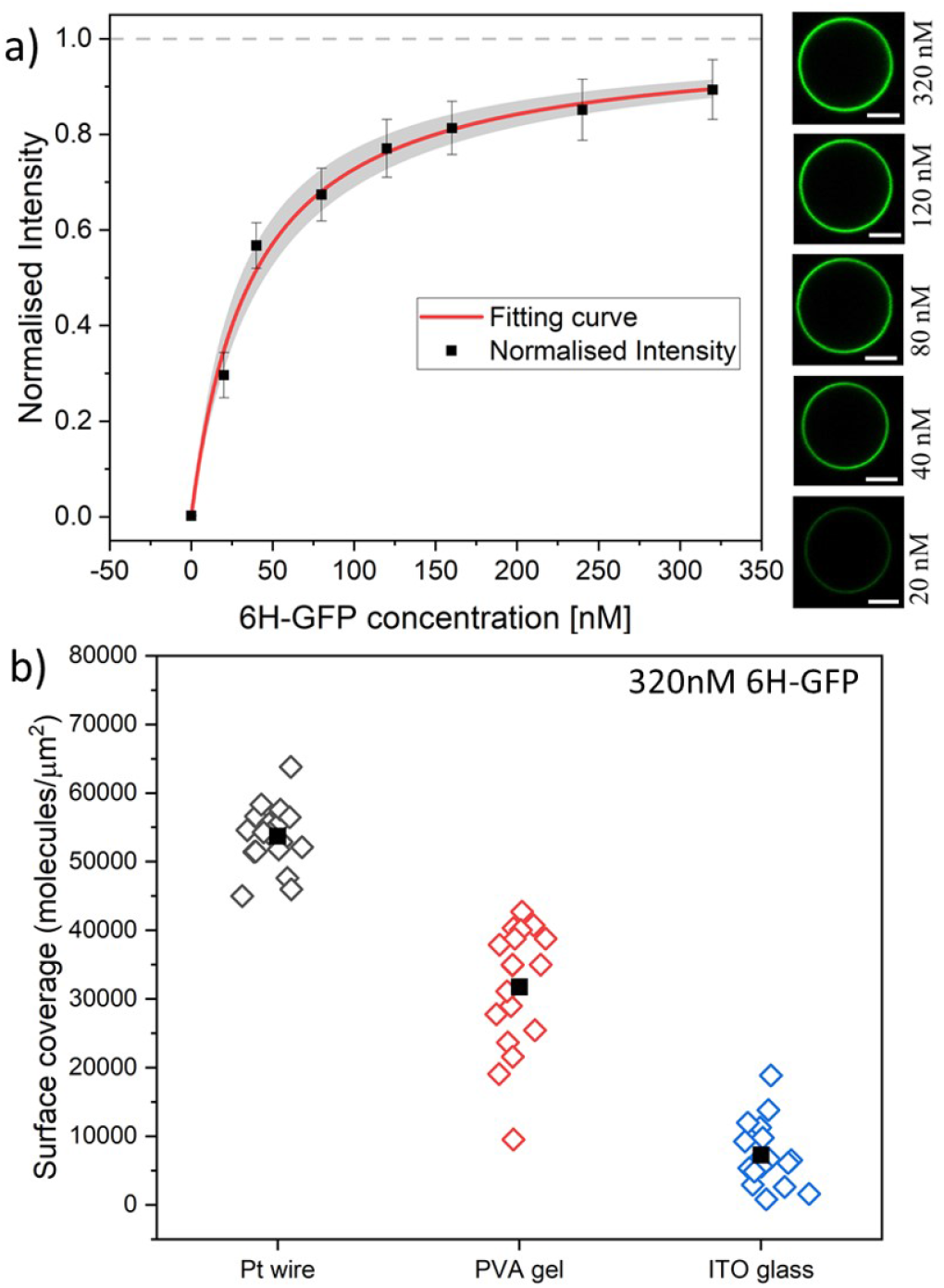
(a) Normalised membrane fluorescence versus molar 6H-GFP concentration, as obtained by the Pt wire protocol. The fitting curve (red) is provided by eqn (6) and leads to the equilibrium dissociation constant *K*_*d*_ = (37.5 ± 7.5) nM for 6H-GFP. The confidence interval for *K*_*d*_ corresponds to the shaded strip (light grey) around the red fitting curve. The images on the right display the brightness of the GUV membranes as directly observed in the microscope for the different molar concentrations. All scale bars: 10 *μ*m; and (b) Surface coverage of GUV membranes by 6H-GFP for different preparation protocols. All GUVs were prepared with 3 mol% of NTA lipids and were exposed to a 320 nM solution of 6H-GFP at pH 7.45. The three sets of data were obtained for 19, 18, and 18 vesicles using the protocol based on Pt wire, PVA gel, and ITO glass, respectively. The solid squares represent the mean values of the coverage obtained for each data set. The numerical values for the mean coverage and the standard deviation of the data in (a) and (b) are given in Tables S1 and S2.

Inspection of Figure 6a shows that the fluorescence intensity becomes saturated for large values of the molar concentration *X*. This saturation regime is characterized by the maximal number of 6H-GFP that can be bound to the NTA anchor lipids. For a lipid bilayer consisting of POPC and cholesterol with a molar ratio of 8:2, the average area per lipid is 0.5 nm^2^ as follows from previous studies ^53,54^. For such a bilayer doped with 3 mol% anchor lipids, neighboring anchor lipids have an average separation of 4.1 nm. This anchor-anchor separation exceeds the lateral size of membrane-bound 6H-GFP, as explained in the *Material and Method* section. Therefore, we conclude that, in the saturation regime, the membrane-bound fluorophores have the same average separation as the anchor lipids, corresponding to a surface coverage of 6.0 × 10^4^ molecules per *μ*m^2^, see eqn (1). In Figure 6b, we use this saturation-based calibration to transform the fluorescence intensities into surface coverages for the three preparation methods.The numerical values of the data in Figures 6a and 6b are given in Tables S1 and S2.

### 2.4 Binding of 6H-FITC to GUV membranes

Next, we studied the His-tagged molecule 6H-FITC, consisting of six histidines attached to the small fluorescent probe FITC. We used the same experimental protocol to determine membrane fluorescence and surface coverage as described in the previous section for 6H-GFP. To obtain the equilibrium dissociation constant for 6H-FITC, we varied the molar concentration *X* of 6H-FITC over the range 0 ≤ *X* ≤ 480 nM and measured the *X*-dependent fluorescence intensity of the GUV membranes, which were prepared by platinum wire electroformation. The corresponding normalised intensity *I*_flu_*/I*_sat_ is plotted in Figure 7a with *I*_sat_ = 40.9 a.u.. These data are well fitted by the intensity-concentration relationship as given by eqn (6) in the *Material and Method* section. From this fit, we obtain the equilibrium dissociation constant *K*_*d*_ = (18.5 ± 3.7) nM for 6H-FITC.

**Fig. 7:**
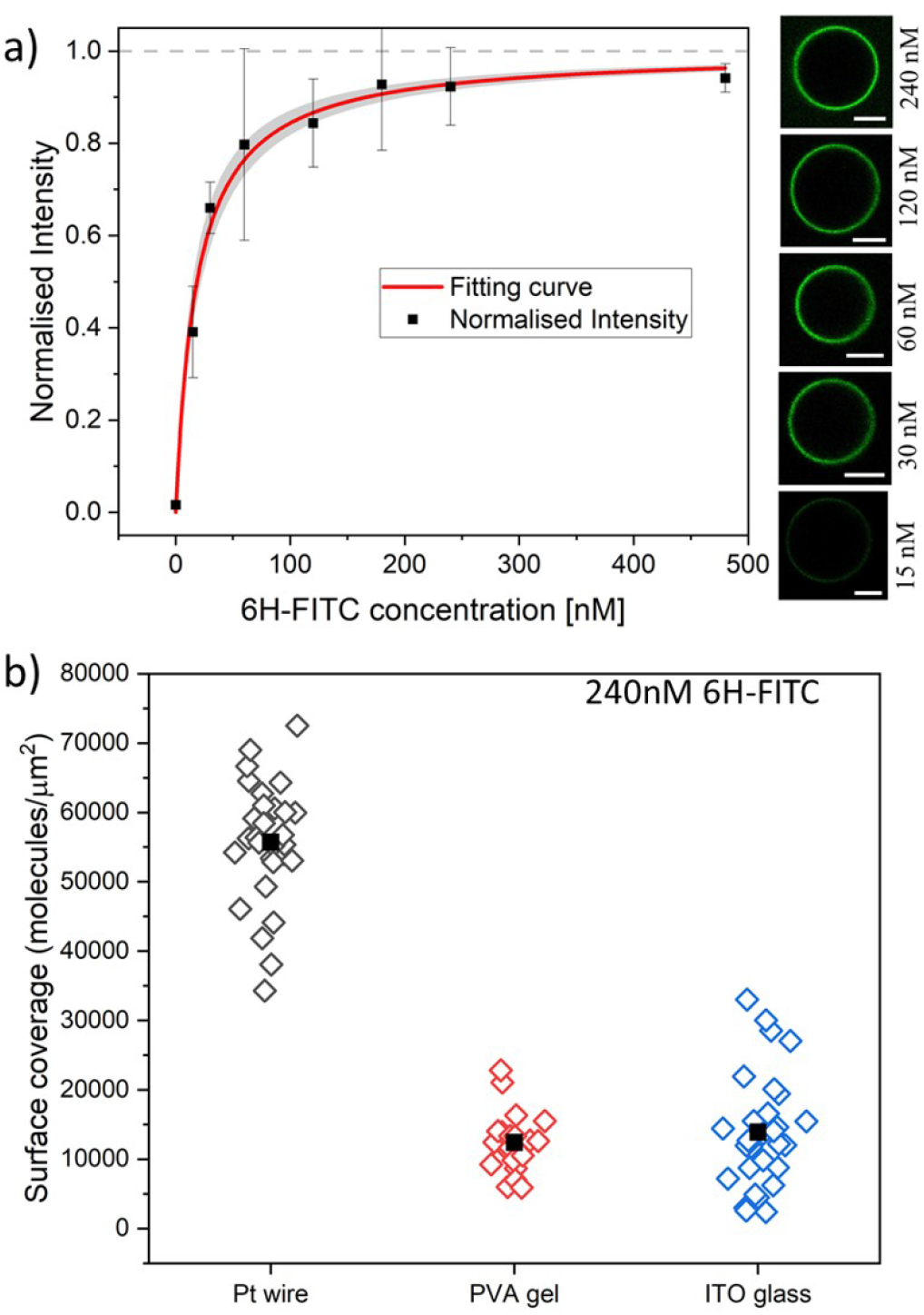
(a) Normalised membrane fluorescence versus molar 6H-FITC concentration, as obtained by the Pt wire protocol. The fitting curve (red) is provided by eqn (6) and leads to the equilibrium dissociation constant *K*_*d*_ = (18.5 ± 3.7) nM for 6H-FITC. The confidence interval for *K*_*d*_ corresponds to the shaded strip (light grey) around the red fitting curve. The images on the right display the brightness of the GUV membranes as directly observed in the microscope for the different molar concentrations. All scale bars: 10 *μ*m; and (b) Surface coverage of GUV membranes by 6H-FITC for different preparation protocols. All GUVs were prepared with 3 mol% of NTA lipids and were exposed to a 240 nM solution of 6H-FITC at pH 7.45. The three sets of data were obtained for 17, 11, and 12 vesicles using the protocol based on Pt wire, PVA gel, and ITO glass, respectively. The solid squares represent the mean values of the coverage for each data set. The numerical values of the mean coverage and the standard deviation of the data in (a) and (b) are provided in Tables S3 and S4.

Comparison of the equilibrium dissociation constant for 6H-FITC with the one for 6H-GFP as given by *K*_*d*_ = 37.5 nM reveals that 6H-FITC is more strongly bound to the GUV membranes compared to 6H-GFP. Furthermore, the different values for the saturation intensity *I*_sat_ imply that the fluorescence of membrane-bound 6H-FITC is strongly reduced compared to the one of membrane-bound 6H-GFP, in agreement with the results on fluorescence quenching by NTA lipids in Figure 5.

The GUV membranes in Figure 7a and Figure 6a contain the same mole fraction of 3 mol% NTA lipids. Therefore, the saturation values of the surface coverage by the two fluorophores are also identical and equal to 6.0 × 10^4^ molecules per *μ*m^2^. In Figure 7b, we use this saturation-based calibration to transform the fluorescence intensities into surface coverages for the three preparation methods. The numerical values corresponding to the data in Figures 7a and 7b are given in Table S3 and S4.

### 2.5 Different compositions of lipid bilayers

#### Dependence on mole fraction of NTA lipids

To get additional insight into the weak binding of His-tagged molecules to GUVs that are grown by electroformation on ITO glass, we compare this method with electroformation on platinum wire, using two different lipid mixtures with mole fractions of 3 mol% and 30 mol% DGS-NTA(Ni) lipids. After harvesting the GUVs, they are exposed to 120 nM 6H-FITC in the exterior solution. As displayed in Figure S2, GUVs prepared by electroformation on ITO glass exhibit a smaller membrane fluorescence than GUVs prepared by electroformation on platinum wire, indicating that only a fraction of the DGS-NTA(Ni) lipid deposited on the ITO glass becomes incorporated into the vesicles. However, for both electroformation methods, the fluorescence increases with increasing mole fraction of the NTA lipids.

A possible explanation for the low membrane fluorescence of GUVs prepared by ITO glass electroformation is as follows. After the deposition of lipid stock on the ITO glass surface, the lipid molecules have orientational freedom to arrange themselves into bilayers ^42^. The surface properties of the ITO glass are likely to affect the properties of the produced vesicles ^55^. The ITO film consists of Indium(III) ions, which can bind to NTA ^56–58^. Presumably, the NTA lipid prefers to stay close to the ITO surface and forms a weak bond with indium, when the lipid stock dissolved in chloroform is deposited on the ITO surface. When the fraction of NTA is increased in the lipid stock to 30 mol%, the ITO surface becomes saturated and some excess NTA is then incorporated into the vesicle membrane.

#### Dependence on mole fraction of cholesterol

GUVs with two different lipid composition are prepared by platinum wire electroformation and exposed to 120 nM 6H-FITC. Both lipid compositions contain 3 mol% NTA lipids but differ in their mole fractions of cholesterol. For lipid bilayers with 20 mol% cholesterol, the intensity is about twice as large as for bilayers without cholesterol, see Figure S3. This change in intensity can arise by two different mechanisms. First, cholesterol may reduce the fluorescence quenching by the NTA lipids; second it may increase the surface coverage by 6H-FITC.

### 2.6 pH dependence of membrane fluorescence

The data on membrane fluorescence and surface coverage as displayed in Figures 6 and 7 are obtained for pH 7.4. These two quantities depend on the pH-value as can be directly concluded from the brightness of the GUV membranes, which varies with the pH, see Figure 8. To determine the pH-dependence of the fluorescence intensity in a systematic manner, the vesicles were grown using platinum wire electroformation. The swelling solution consisted of 50 mM sucrose and 2 mM HEPES buffer at pH 7.4. The vesicles were then exposed to an exterior solution containing 20 nM 6H-GFP or 120 nM 6H-FITC as well as 22.5 mM NaCl, 5 mM sucrose and 2 mM HEPES buffer. The pH of these solutions was increased and decreased by adding NaOH and HCl, respectively.

**Fig. 8:**
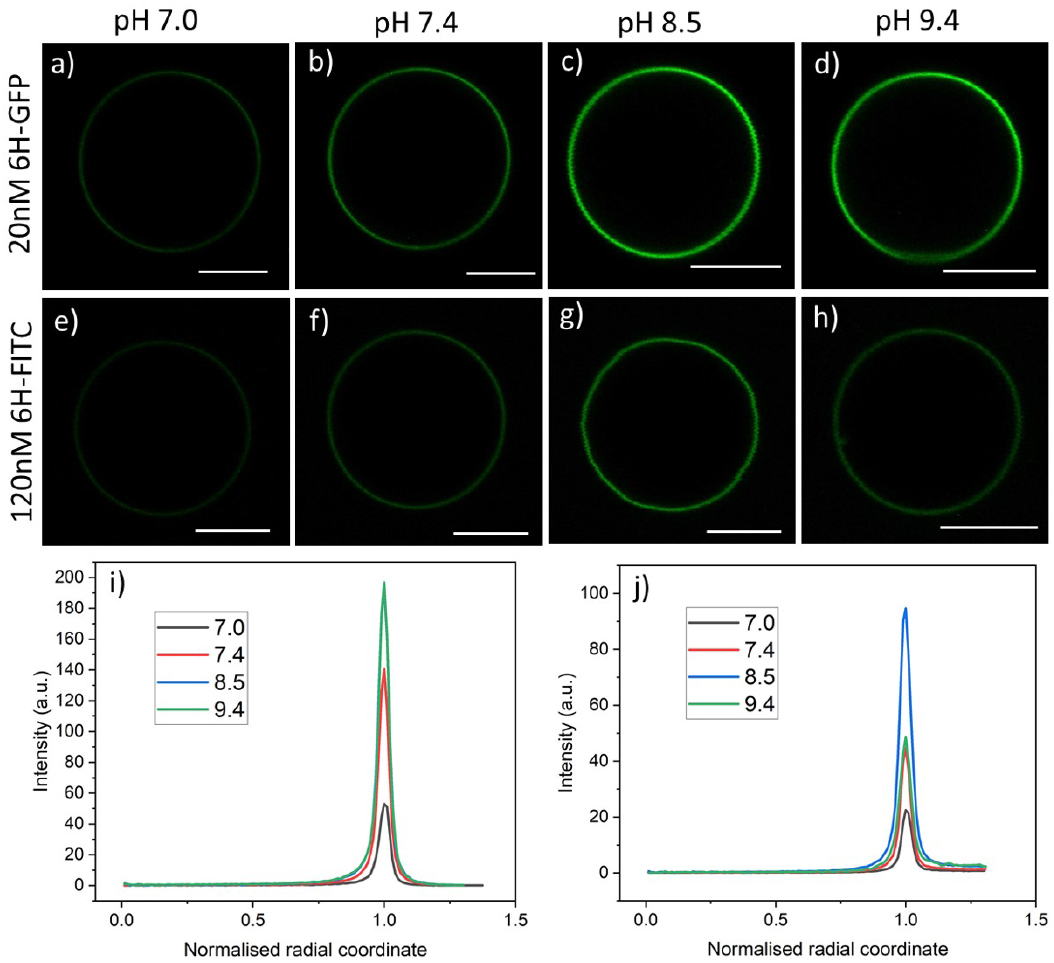
Brightness of membrane fluorescence for 3 mol% NTA lipids and two types of His-tagged molecules, 6H-GFP and 6H-FITC, as observed for different pH-values of the exterior solution: GUVs exposed to 20 nM solution of 6H-GFP in (a-d) and to 120 nM of 6H-FITC in (e-h). All GUVs are prepared using the platinum wire method. All scale bars: 10 *μ*m; and (i,j) Corresponding fluorescence intensity profiles as obtained by the method described in Figure 4. The intensity profiles in (i) and (j) correspond to the images in (a-d) and (e-h), respectively.

For each pH value, the membrane fluorescence was determined via the method described in Figure 4. The resulting fluorescence intensities of the GUV membranes are displayed in Figure 9a and 9b for 6H-GFP and 6H-FITC, respectively. For 6H-GFP, the membrane fluorescence first increases and then saturates for pH values above pH 8.5. For 6H-FITC, the intensity exhibits a pronounced maximum at about pH 8.5. The numerical values of the data in Figures 9a and 9b are given in Tables S5 and S6.

**Fig. 9:**
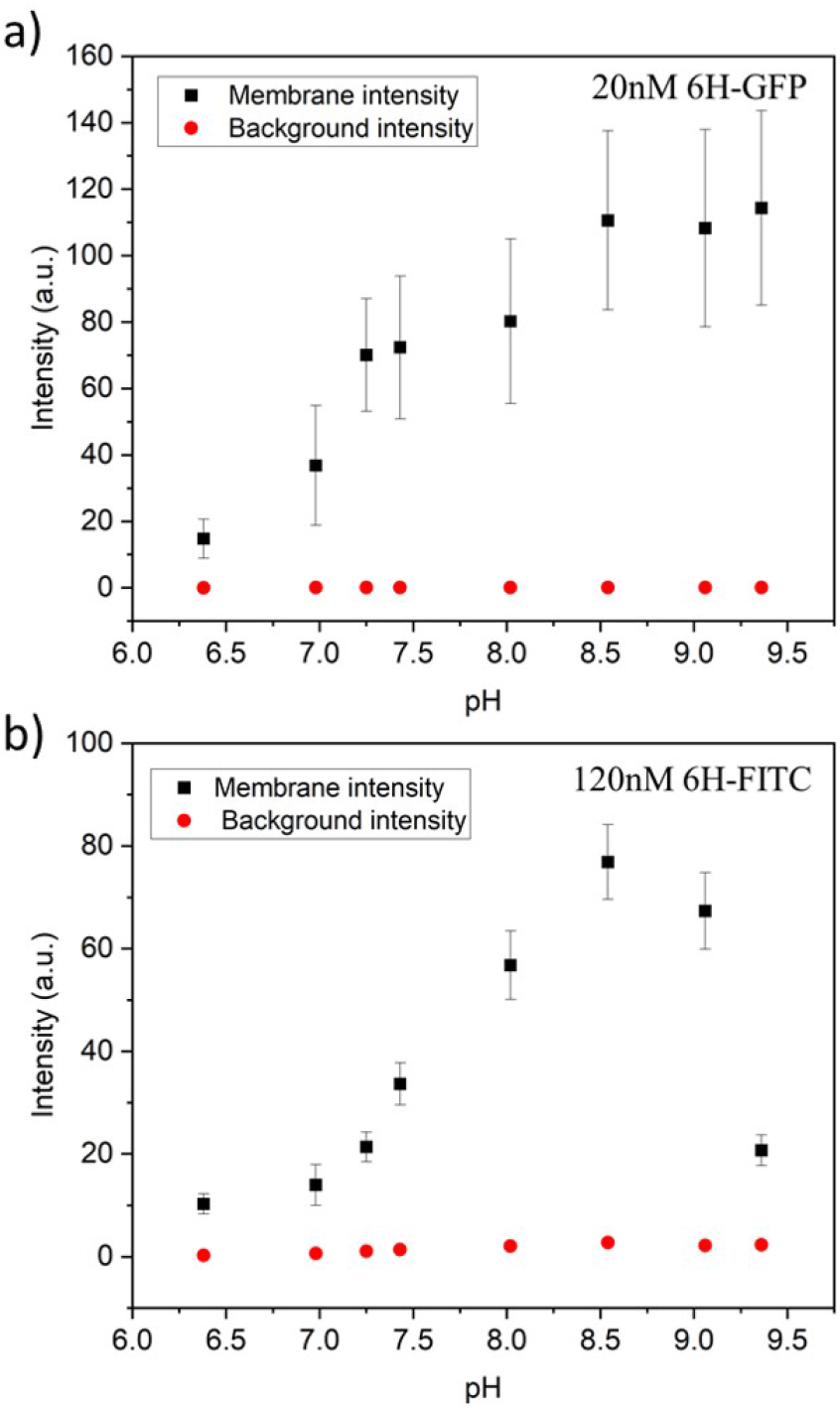
Membrane intensities (black squares) and background intensities (red circles) versus pH of exterior solution for GUV membranes with 3 mol% NTA lipids. All GUVs were prepared by electroformation on platinum wire: (a) For GUVs exposed to 20 nM 6H-GFP, the membrane intensity increases with increasing pH for 6.38 ≤ pH ≤ 8.5 and then saturates; and (b) For GUVs exposed to 120 nM 6H-FITC, the membrane intensity exhibits a pronounced maximum close to pH 8.5. The numerical values of the data in (a) and (b) are given in Tables S5 and S6. For both 6H-GFP and 6H-FITC, the background intensities (red circles) are negligible over the whole range of pH values, which demonstrates that the pH dependence of the membrane fluorescence arises from the pH dependence of the His-NTA binding affinity.

In general, the fluorescence of a dye molecule can vary with the pH value even in the absence of lipids. In order to determine the latter pH dependence, we also measured the background intensities for 6H-GFP and 6H-FITC in the exterior solution far away from the vesicle membranes, see the corresponding data in Figure 9. Inspection of both panels shows that the background intensity in the exterior solution is negligible over the whole range of pH values studied here. Therefore, the pH dependence of the membrane fluorescence in Figure 9 is caused by the pH dependence of the membrane-bound fluorophores rather than by the intrinsic pH dependence of these dye molecules in solution.

## 3 Conclusions and Outlook

The binding of His-tagged molecules to GUV membranes has been investigated, using the anchor lipids DGS-NTA(Ni) that form coordinate bonds with the His-tags (Figure 2). Two different green fluorescent molecules, 6H-GFP and 6H-FITC, have been studied, both of which were tagged to a chain of six histidines. Using three different preparation methods - electroformation on platinum wire, swelling of PVA hydrogels, and electoformation on ITO glass - as well as microfluidics (Figure 10) to expose the GUVs to a constant bulk concentration of the fluorophores, we obtained the images in Figure 3 which show that the brightness of the GUV membranes depends both on the type of His-tagged molecule and on the preparation method. To determine the membrane fluorescence in a quantitative manner, we measured the fluorescence intensity profiles, see Figure 4a, and obtained the membrane fluorescence by integrating the excess intensity profiles as in Figure 4b over the radial coordinate.

**Fig. 10:**
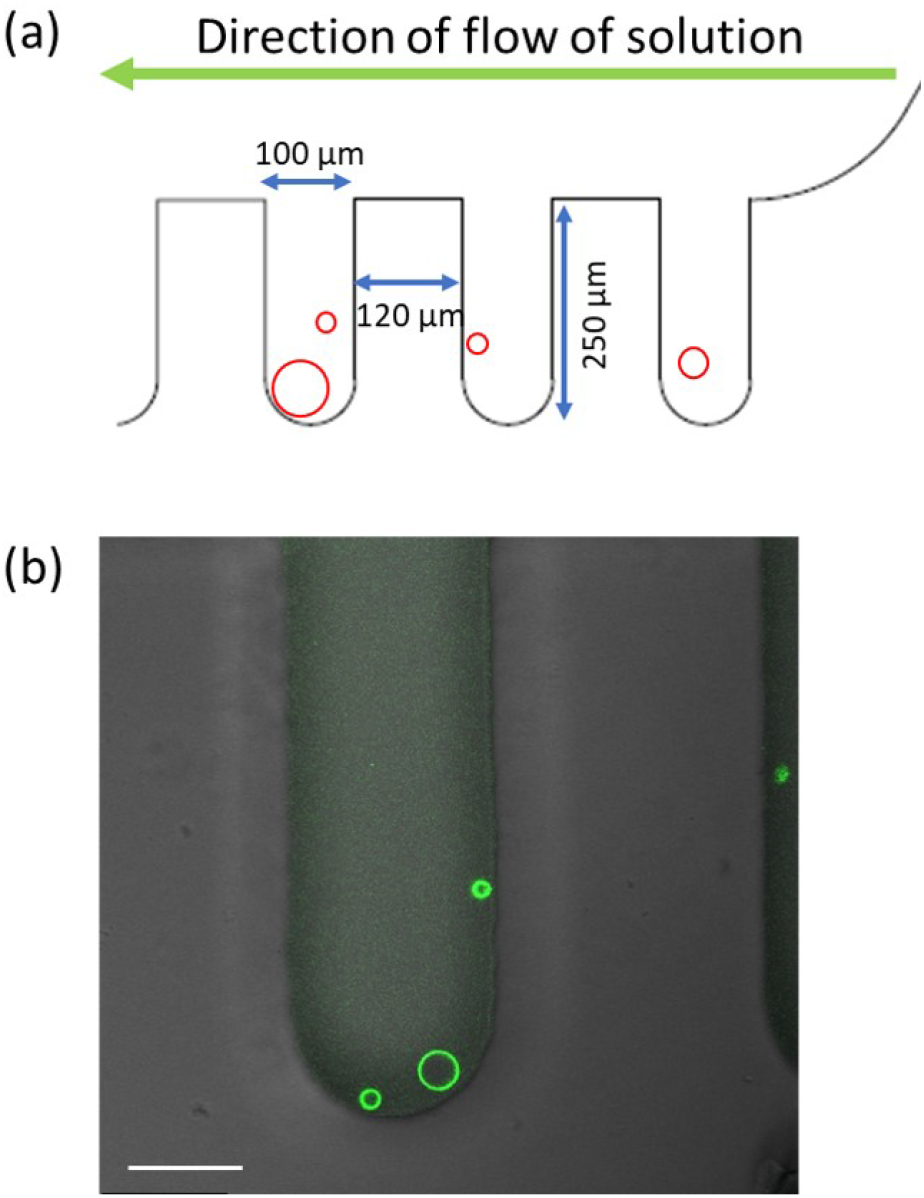
Microfluidic approach used to control the concentration of His-tagged molecules in the exterior solution of the GUVs: (a) Design of chip with dead-end side channels perpendicular to the main channel and the direction of flow. The His-tagged molecules have a constant concentration in the main channel and reach the GUVs in the side channels only by diffusion; and (b) Overlay of fluorescence and differential interference contrast microscopy for one side channel with three trapped GUVs exposed to 640 nM 6H-GFP. Scale bar: 50 *μ*m.

One surprising result of our study is the different behavior of the two fluorophores when exposed to an increasing amount of NTA lipids. Indeed, we observed essentially no change in the fluorescence of 6H-GFP, see Figure 5a, but a strong quenching effect on 6H-FITC as demonstrated by the data in Figure 5b.

Both for 6H-GFP and 6H-FITC, the highest brightness of the GUV membranes was obtained when we prepared these vesicles via electroformtion on platinum wire. We used this latter method to measure the dependence of the fluorescence intensities on the molar concentrations of the His-tagged fluorophores (Figures 6a and 7a). Both sets of data are well fitted by the intensity-concentration relationship in eqn (6), which depends on only two parameters, the equilibrium dissociation constant *K*_*d*_ and the saturation intensity *I*_sat_.

The equilibrium dissociation constant turns out to be *K*_*d*_ = (37.5 ± 7.5) nM for 6H-GFP and *K*_*d*_ = (18.5 ± 3.7) nM for 6H-FITC which implies that 6H-FITC is more strongly bound to the GUV membranes. On the other hand, the strong quenching of 6H-FITC by the Ni^2+^ ions of the NTA lipids (Figure 5b) leads to the saturation intensity *I*_sat_ = 40.9 a.u. for 6H-FITC which is much smaller than *I*_sat_ = 320.4 a.u. for the membrane fluorescence of 6H-GFP.

Our method to determine the equilibrium dissociation constant *K*_*d*_ is quite general and can be applied to the binding between other His-tagged fluorophores and/or other anchor lipids. First, one measures the fluorescence intensities as a function of the molar concentration *X* and then plots the normalised fluorescence intensities *I*_flu_*/I*_sat_ versus *X* as in Figures 6a and 7a. Second, one fits the plotted data by the intensity-concentration relationship in eqn (6), which involves two fit parameters, the equilibrium dissociation constant *K*_*d*_ and the saturation intensity *I*_sat_. If the data can be well-fitted by this equation, one obtains a reliable estimate for *K*_*d*_. On the other hand, if the fit turns out to be unacceptable, one can conclude that the model used to derive eqn (6) should be modified.

The most important simplifying assumption of this model is that each anchor lipid can bind one fluorophore. Therefore, a poor fit will provide direct evidence that the latter assumption does not apply to the system under consideration. Possible modifications of the model include (i) crowding, i.e., the steric hindrance between the membrane-bound fluorophores, (ii) the binding of several fluorophores to one anchor lipid, and (iii) the recruitment of several anchor lipids by one fluorophore.

Another surprising outcome of our study is that it reveals a pronounced pH dependence of the membrane fluorescence for both 6H-GFP and 6H-FITC, see Figure 9a and b. For 6H-GFP, the membrane fluorescence first increases and then saturates for pH values above pH 8.5. For 6H-FITC, on the other hand, the membrane fluorescence exhibits a pronounced maximum at about pH 8.5 (Figure 9b). Furthermore, the observed membrane fluorescence was always much larger than the background intensity, corresponding to those fluorophores in the exterior solution that were not in contact with the lipid membranes. Therefore, the pH dependence displayed in Figures 9 represents the the pH dependence of the membrane fluorescence rather than the intrinsic pH dependence of the fluorescent molecules in solution.

Previous experiments have shown that GUV membranes exposed to nanomolar solutions of 6H-GFP aquire a large spontaneous curvature that can be fine-tuned to divide the GUVs in a controlled manner. ^28^ As shown here, 6H-FITC has a smaller equilibrium constant and is, thus, more strongly bound to GUV membranes compared to 6H-GFP. Therefore, it will be interesting to study the spontaneous curvature generated by 6H-FITC and the shape transformations of GUVs arising from this curvature generation.

## 4 Material and Method

### 4.1 Preparation of giant unilamellar vesicles

Chloroform stock solutions of 1-palmitoyl-2-oleoyl-sn-glycero-3-phosphocholine (POPC) and cholesterol were mixed at a molar ratio of 8:2 with a final lipid concentration of 4 mM. The required mol% of DGS-NTA (Ni) was added to the lipid stock solution. This stock was used as it is or diluted depending on the preparation method. All lipids were obtained from Avanti Polar Lipids.

His-tagged GFP was obtained from Astrid Krämer of the protein facility at the Max Planck Institute of Molecular Physiology (Dortmund, Germany). His-tagged FITC was purchased from Biomatik Corporation (Ontario, Canada).

#### Electroformation on platinum wire ^32,41^

6 *μ*L of a 0.5 mM solution of the same lipid mixture was spread on two platinum wires using a syringe. The wires were kept under vacuum for an hour to remove traces of chloroform. The wires were then dipped in a quartz cuvette, which was filled with 50 mM of sucrose in 2 mM HEPES buffer (pH 7.4). An AC electric field with peak-to-peak voltage of 3V and frequency of 10 Hz was applied for 2 hours at 35°C to speed up the vesicle swelling process. Subsequently, the frequency of the AC field was reduced to 3Hz and the low-frequency field was applied for 5 min, to detach the vesicles from the wires. After cooling to room temperature, the vesicles were harvested using a micropipette with a broad tip.

#### PVA hydrogel swelling. ^28,36,39^

A PVA solution was prepared by dissolving 40 mg of PVA in 1 mL of water. 40 *μ*L of this solution was spread on clean cover glasses and dried in an oven at 40°C for 30 min. 4 *μ*L of 2 mM lipid mixture consisting of POPC, cholesterol, and NTA was spread and kept under vacuum for one hour to eliminate trace amounts of chloroform. A closed chamber was created using a Teflon spacer and another cover glass. An aqueous solution of 50 mM sucrose in 2 mM HEPES buffer (pH 7.4) was introduced into the chamber to hydrate the lipid film at room temperature (23°C). The vesicles were harvested after gentle tapping on the PVA coated glass.

#### Electroformation on ITO glasses. ^42–44^

Commercially available ITO glass from PGO-GmbH (Iserlohn, Germany) was used without further treatment. In general, the ITO glass surface can have a complex molecular structure, depending on the pretreatment conditions. ^59^ 4 *μ*L of a 4 mM solution of the lipid mixture was spread on two ITO coated glasses using a syringe. The resulting lipid films formed bilayers stacks on the ITO glass surfaces, which were then kept under vacuum for an hour to remove remaining traces of chloroform. The two glasses were then arranged into a closed chamber by using a Teflon spacer as side walls. The chamber was filled with 50 mM of sucrose in 2 mM HEPES buffer (pH 7.4). An AC electric field with peak-to-peak voltage of 2V and frequency of 10 Hz was applied for 2 hours at room temperature (23°C). Subsequently, the frequency of the AC field was reduced to 3Hz and the low-frequency field was applied for 5 min, to detach the vesicles from the glass surfaces. The vesicles were harvested using a micropipette with a broad tip.

### 4.2 Microfluidic observation chamber

The experiments were performed on polydimethylsiloxane (PDMS) microfluidic chips. The design of the device is such that the vesicles are trapped in the dead-end side channels that are perpendicular to the main channel, as in Figure 10a. The global design of the microfluidic chip is displayed in Figure S4. The solution volume that can be loaded in the microfluidic device is 2.5 *μ*L and each of the side channels is 250 *μ*m deep and 100 *μ*m wide. The reservoir in the inlet is filled with vesicles and NaCl solution. The vesicles are loaded on the chip using a syringe connected to a pump at the outlet. Then the chip is oriented vertically in such a way that the vesicles slowly fall into the side channels. After about 20 mins, the chip is placed on the microscope. The solution in the reservoir is exchanged by another solution with a certain concentration of HIs-tagged molecules. Because of the continuous flow from the inlet, the concentration of these molecules remains constant during the whole experiment.

On the other hand, the His-tagged molecules reach the vesicles in the side channels only by diffusion which implies that the vesicles are shielded from the hydrodynamic flow and do not suffer any mechanical perturbations that could otherwise arise from this flow. The trapping in the side channels also helps to monitor the same GUV throughout the experiment. As shown in Movie 1, the complete solution exchange including the side channels takes around 3.5 mins. For all experiments performed here, the pump was used to exchange 20 *μ*L of the his-tagged molecules, corresponding to 8 times the solution volume within the microfluidic device, before the vesicles have been imaged.

### 4.3 Quantum yield of fluorophores in the absence of lipids

To determine the different quantum yields of the two fluorophores, the detector gain was adjusted to give the same intensity for the same molar concentrations of 6H-FITC and 6H-GFP in the absence of lipids. Briefly, 100 nM solutions of 6H-FITC and 6H-GFP were imaged using the HyD detector on the Leica SP8 confocal microscope. Both molecules were excited using a 488 nm Argon laser and emission was collected between 495 and 550 nm. The detector gain was adjusted to obtain the same brightness for both solutions.

From this adjustment, we concluded that 6H-FITC is about 1.2 times brighter than 6H-GFP.

### 4.4 Measurements of fluorescence quenching

Experiments to observe the possible quenching of the fluorescence of the His-tagged fluorophores by the DGS-NTA(Ni) lipids were conducted using the FluorMax-4 fluorimeter. LUVs were prepared by hydrating dried lipids films (POPC: Cholesterol (8:2) and 30 mol% DGS-NTA(Ni)) with 2mM HEPES buffer. After vortexing for 30 mins, the suspension was extruded 21 times through a 200nm polycarbonate filter (Avanti mini extruder). Fluorescence spectra were recorded for 120nM 6H-GFP and 120nM 6H-FITC in the presence and in the absence of the LUVs by exciting at 488nm. Control experiments were also performed for LUVs without the DGS-NTA(Ni) lipids, see Figure S1.

### 4.5 Image acquisition

All fluorescent images were recorded using a Leica SP8 confocal microscope. Argon laser with wavelength 488 nm was used to excite both the GFP proteins and the FITC dye. The emission was collected in the range of wavelengths from 495 to 550 nm. The images for quantification of the fluorescence intensity on the membrane were obtained using a 40X 1.3 oil immersion objective.

### 4.6 Separation of NTA lipids and lateral size of fluorophores

For a lipid bilayer consisting of POPC and cholesterol with a molar ratio of 8:2, the average area per lipid is 0.5 nm^2^ as follows from previous studies ^53,54^. When such a bilayer is doped with 3 mol% NTA lipids, these anchor lipids have an average spatial separation of 4.08 nm. This separation determines the surface density *ρ*_anc_ of the NTA lipids via

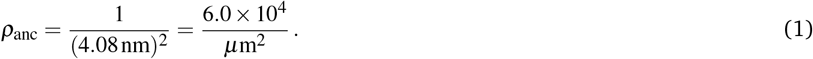

When each NTA anchor lipid is occupied by one His-tagged molecule, corresponding to the saturation regime, the surface density *ρ*_anc_ of the anchor lipids is equal to the surface density or coverage of the His-tagged fluorophores.

In order to estimate whether or not this saturation regime leads to crowding of the membrane-bound fluorophores, we need to compare the average separation of the NTA lipids with the lateral size of the membrane-bound fluorophores. The lateral size of membrane-bound 6H-GFP should be comparable with the diameter of its *β*-barrel. This diameter is about 3 nm ^28,60^ which is smaller than the anchor-anchor separation of 4.08 nm and implies that crowding effects between membrane-bound 6H-GFP molecules can be ignored even when each NTA anchor lipid is occupied by one such molecule. The 6H-FITC peptide, on the other hand, has a linear extension of about 3.2 nm which provides an upper bound on the lateral size of membrane-bound 6H-FITC. Because this upper bound on the lateral size is again smaller than the average separation of the NTA anchor lipids, crowding effects between membrane-bound 6H-FITC molecules can again be ignored even when each NTA lipid binds one such molecule.

### 4.7 Fluorescence of membrane-bound fluorophores

Radial Profile Angle from ImageJ was used to plot the fluorescence intensity as a function of the radial coordinate *r*, see Figure 4a. The peak analyzer of Origin2021b from OriginLab was used to determine the excess intensity in Figure 4b by subtracting the baseline intensity from the fluorescence intensity. Then the membrane fliuorescence was obtained by integrating the excess intensity over the radial coordinate *r*, using the whole range of *r*-values.

To measure the membrane fluorescence at different pH values, the vesicles were grown using platinum wire electroformation at pH 7.4. These vesicles were then trapped and exposed to the solution of the His-tagged molecules at different pH, ranging from 6.4 to 9.4. The pH of the solution was adjusted using HCl or NaOH. The final pH of the solutions was measured using a micro pH electrode from Mettler Toledo. The results of these measurements are displayed in Figure 9.

### 4.8 Binding kinetics and equilibrium dissociation constant

The binding sites for the His-tagged molecules are provided by the NTA anchor lipids. We consider a simple binding model and assume that each anchor lipid can bind at most one His-tagged molecule. The outer leaflet of the GUV membrane contains a total number of *N*_anc_ anchor lipids which can bind *N*_bH_ His-tagged molecules with *N*_bH_ ≤ *N*_anc_. For each anchor lipid, the binding (or association) rate is taken to be proportional to the molar concentration *X* and, thus, has the form *κ*_on_ *X* with the association rate constant *κ*_on_, while unbinding (or dissociation) is an activated process with the dissociation rate *ω*_off_. The number *N*_bH_ of bound His-tagged molecules then changes with time *t* according to the evolution equation

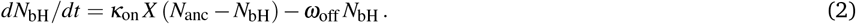

In equilibrium, the binding flux is equal to the unbinding flux which implies *dN*_bH_*/dt* = 0. The equilibrium value 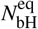 of the bound His-tagged molecules is then given by

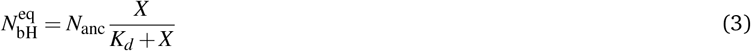

which depends on the equilibrium dissociation constant

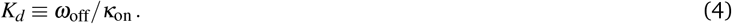

When we divide eqn·(3) by the surface area *A* of the membrane, we obtain the equilibrium surface density or coverage 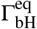 of the His-tagged molecules which depends on the surface density *ρ*_anc_ of the NTA anchor lipids and on the molar concentration *X* according to

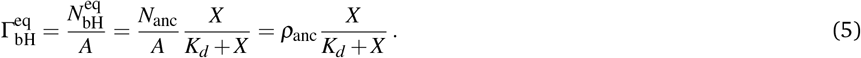

For 3 mol% NTA lipids, the surface density *ρ*_anc_ is equal to 6.0 × 10^4^ molecules per *μ*m^2^ as in eqn (1).

### 4.9 Fluorescence intensity of membrane-bound fluorophores

The fluorescence intensity of the membrane-bound fluorophores is proportional to their surface density. Because the fluorescence of these fluorophores can be quenched by the NTA anchor lipids, we introduce a reduction factor *ϕ* with 0 *< ϕ* ≤ 1 for the fluorescence of a fluorophore bound to the NTA lipid. The data in Figure 5 imply that *ϕ* = 1 for 6H-GFP and *ϕ <* 1 for 6H-FITC. We then obtain the relationship

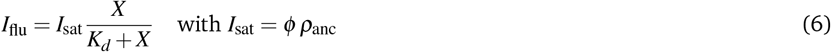

between the fluorescence intensity *I*_flu_ and the molar concentration *X*. As a consequence, the fluorescence intensity *I*_flu_ increases monotonically with the molar concentration *X* and attains its saturation value *I*_flu_ ≈ *I*_sat_ = *ϕρ*_anc_ for large *X* as in Figures 6a and 7a.

For 6H-GFP and 6H-FITC, the fluorescence intensities attain the saturation values *I*_GFP_ = 320.4 a.u. and *I*_FITC_ = 40.9 a.u. as obtained from the HyD detector on the Leica SP8 confocal microscope. Taking into account that the data in Figures 6b and 7b were obtained for GUV membranes with the same mole fraction of 3 mol% NTA lipids and thus with the same surface density *ρ*_anc_, we obtain the ratio

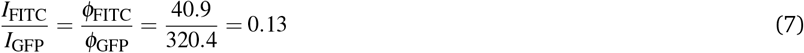

of the two fluorescence reduction factors *ϕ*_FITC_ and *ϕ*_GFP_. It then follows from *ϕ*_GFP_ = 1 for 6H-GFP, that the fluorescence of membrane-bound 6H-FITC molecules is reduced by the factor *ϕ*_FITC_ = 0.12.

### 4.10 Calibration of surface coverages via saturation regime

The saturation regime of the His-tagged fluorophores, which corresponds to the limit of large molar concentrations *X*, is used to calibrate the transformation from membrane fluorescence to surface coverage. For large *X*, the coverage 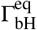 of the membrane-bound fluorophores is equal to the surface density *ρ*_anc_ of the NTA anchor lipids as in eqn (5). A combination of this latter equation with eqn (6) leads to

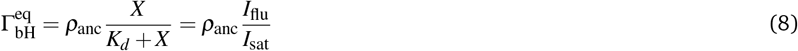

which allows us to compute the surface coverage 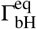 of the His-tagged molecules from (i) the surface density *ρ*_anc_ of the NTA anchor lipids, see eqn (1), and (ii) the data for the normalized fluorescence intensity *I*_flu_*/I*_sat_ as displayed in Figures 6a and 7a.

## Abbreviations

AC: alternating current
DGS-NTA(Ni) or NTA for short: 1,2-dioleoyl-sn-glycero-3-[(N-(5-amino-1-carbox-ypentyl)iminodiacetic acid)succinyl] (nickel salt)
FITC: Fluorescein isothiocyanate
GFP: green fluoerescent protein
ITO: indium tin oxide
POPC: 1-Palmitoyl-2-oleoyl-sn-glycero-3-phosphocholine
PVA: polyvinyl alcohol
6H-GFP: green fluorescent protein tagged by six histidines
6H-FITC: Fluorescein isothiocyanate tagged by six histidines

## Acknowledgements

We acknowledge support by the Max Planck School ‘Matter to Life’. We thank Astrid Krämer for providing the 6H-GFP molecules as well as Tom Robinson and Naresh Yandrapalli for help with the initial setup of the microfluidics.

## Footnotes

* For chemical formulas, see the list of abbreviations at the end of this article.

