## Supporting information for "Binding of His-tagged fluorophores to lipid bilayers of giant vesicles†"

**Electronic Supplementary Information (ESI)**

[a] *Theory and Bio-Systems, Max Planck Institute of Colloids and Interfaces, Science Park Golm, 14424 Potsdam, Germany*

[b] *Center for Synthetic Biology, Northwestern University, Evanston, USA*

This Electronic Supplementary Information contains four Supplementary Figures and seven Supplementary Tables as well as the caption of Movie 1:

**Supplementary Figures:**

**Figure S1**: No fluorescence quenching of 6H-FITC in the absence of NTA lipids

**Figure S2:** Dependence of membrane fluorescence on mole fraction of NTA lipid

**Figure S3:** Dependence of membrane fluorescence on mole fraction of cholesterol

**Figure S4:** Global design of microfluidic chip

**Supplementary Tables:**

**Table S1:** Membrane fluorescence versus molar concentration of 6H-GFP (Figure 6a)

**Table S2:** Surface coverage of GUV membranes by 6H-GFP (Figure 6b)

**Table S3:** Membrane fluorescence versus molar concentration of 6H-FITC (Figure 7a)

**Table S4:** Surface coverage of GUV membranes by 6H-FITC (Figure 7b).

**Table S5:** Membrane fluorescence intensity of 6H-GFP versus pH (Figure 9a)

**Table S6:** fluorescence of membrane-bound 6H-FITC versus pH (Figure 9b).

**Table S7:** Membrane fluorescence for increased mole fractions of NTA (Figure S2).

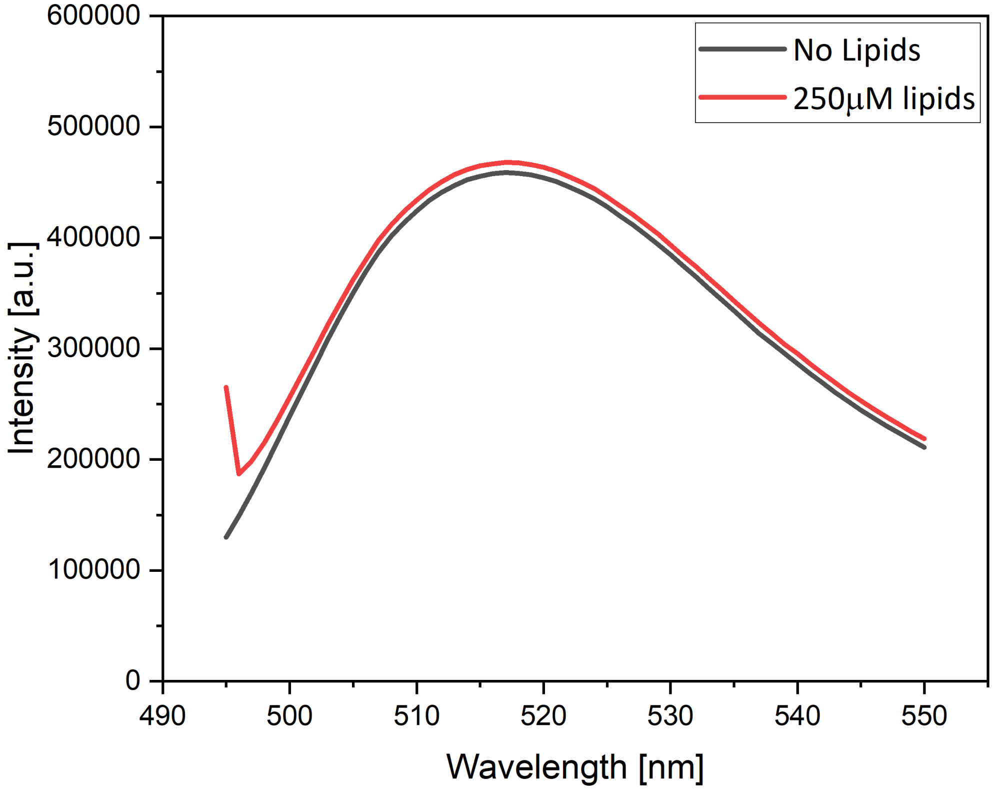


Figure S1: Control experiments to demonstrate that the fluorescence of 6H-FITC is not quenched by liposomes which contain a mixture of POPC and cholesterol with a molar ratio of 8:2 but no NTA lipids.

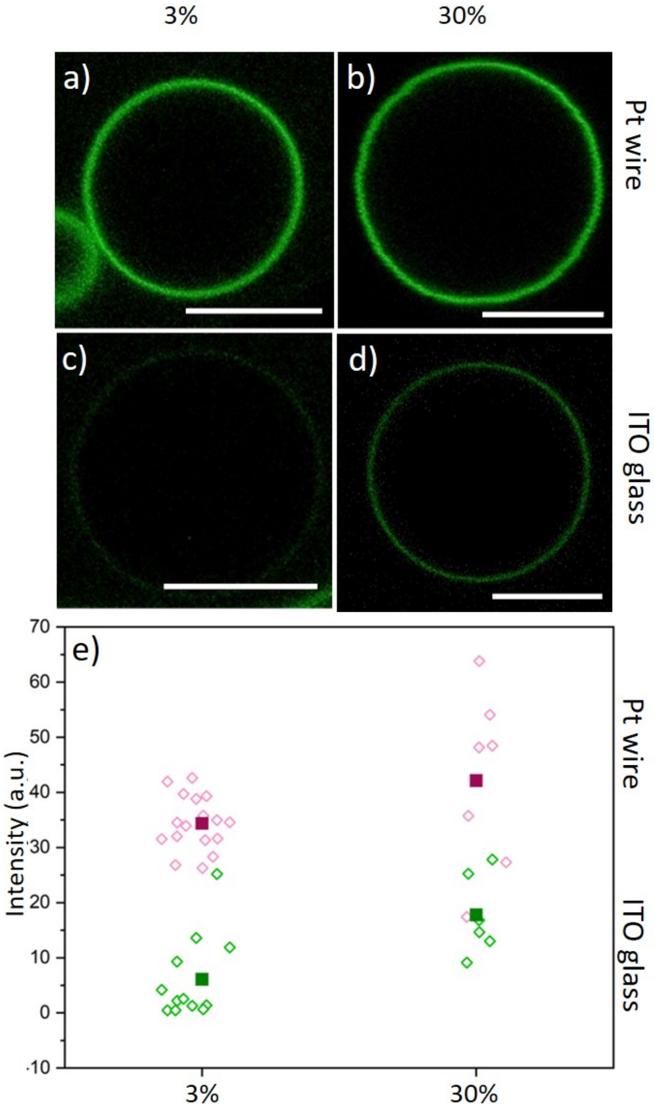


Figure S2: Binding of 6H-FITC to GUV membranes with 3 mol% and 30 mol% DGS-NTA(Ni) anchor lipid: GUVs formed by (a,b) Pt wire electroformation and by (c,d) ITO glass electroformation. All GUVs are exposed to a 120 nM solution of 6H-FITC; and (e) Membrane fluorescence for the four preparation conditions in (a-d). The filled squares represent the mean values of the respective data. The numerical values of these data are given in Table S3. All scale bars are 10 $\mu$m.

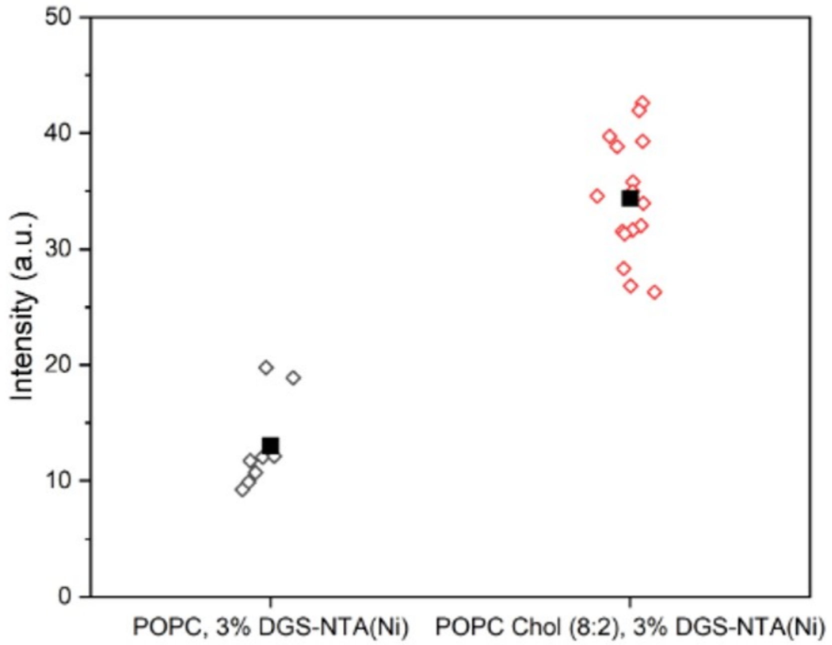


Figure S3: Influence of cholesterol on the fluorescence intensity of GUV membranes: The GUVs are prepared by platinum wire electroformation and exposed to 120 nM 6H-FITC for two different lipid compositions. In the presence of 20 mol% cholesterol, the intensity is about twice as large as in the absence of cholesterol which indicates that cholesterol reduces the fluorescence quenching by the NTA lipids.

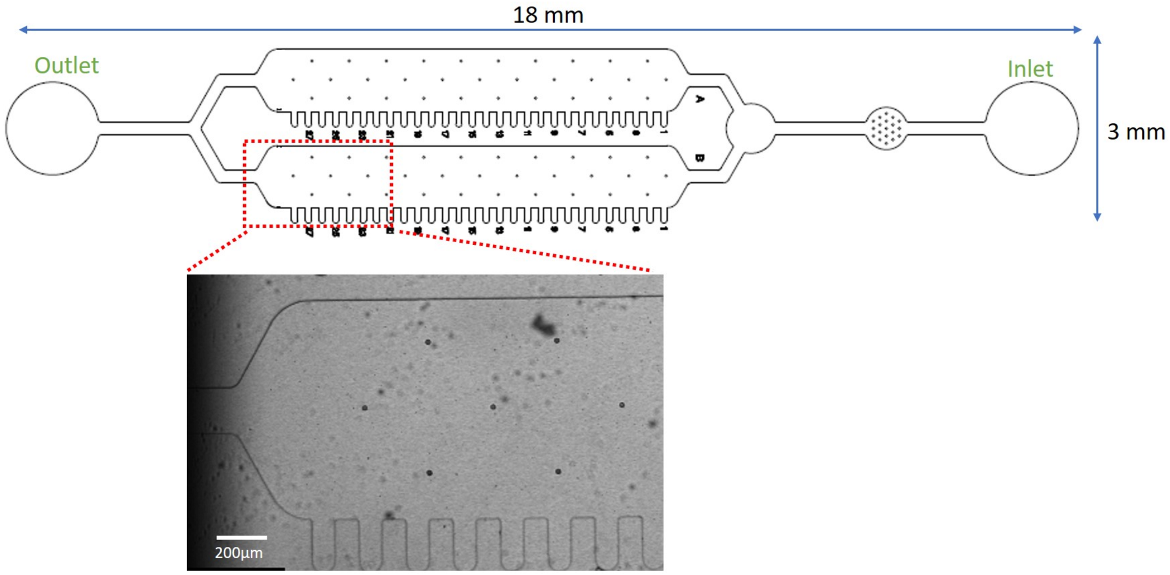


Figure S4: (Top) Design of microfluidic chip: The device has two main channels parallel to each other. A pipette tip is used as the reservoir for the solution at the inlet. A syringe pump connected to the outlet draws the solution from the reservoir to the outlet via the main channels. The side channels trap the GUVs; and (Bottom) Bright field image showing and magnifying the portion of the chip within the broken red frame. The solution within the side channels is exchanged by diffusion from the main channel. The complete solution exchange including the side channels takes about 3.5 mins as shown in Movie 1.

Table S1: Statistics for the normalised fluorescence intensity of GUV membranes exposed to different molar concentrations $X$ of 6H-GFP, as obtained via the platinum wire method, corresponding to the data in Figure 6a. The membranes contained 3 mol% NTA lipids and were exposed to a 320 nM solution of 6H-GFP at pH 7.45. The number of GUVs used for the statistics is denoted by #GUVs and displayed in the 2nd column. The numerical values for the mean value and the standard deviation (SD) of the normalised intensity are dimensionless.

| $X$ [nM] | #GUVs | Mean intensity | SD |
|---|---|---|---|
| 0 | 4 | 0.002 | 0.001 |
| 20 | 4 | 0.296 | 0.047 |
| 40 | 4 | 0.567 | 0.048 |
| 80 | 4 | 0.674 | 0.055 |
| 120 | 4 | 0.771 | 0.061 |
| 160 | 4 | 0.813 | 0.056 |
| 240 | 4 | 0.851 | 0.064 |
| 320 | 4 | 0.894 | 0.062 |

Table S2: Statistics for the surface coverage of GUV membranes by 6H-GFP as obtained for three different preparation methods, corresponding to the data in Figure 6b. The membranes contained 3 mol% NTA lipids and were exposed to a 320 nM solution of 6H-GFP at pH 7.45. The number of GUVs used for the statistics is denoted by #GUVs and displayed in the 2nd column. The numerical values for the mean value and the standard deviation (SD) of the coverage are given in units of membrane-bound molecules per $\mu m^2$.

| Preparation method | #GUVs | Mean coverage | SD |
|---|---|---|---|
| Pt wire electroformation | 19 | 53 703 | 4 449 |
| PVA hydrogel swelling | 18 | 31 742 | 9 071 |
| ITO glass electroformation | 18 | 7 261 | 4 582 |

Table S3: Statistics for the normalised fluorescence intensity of GUV membranes exposed to different molar concentrations $X$ of 6H-FITC, as obtained by the platinum wire method, corresponding to the data in Figure 7a. The membranes contained 3 mol% NTA lipids and were exposed to a 240 nM solution of 6H-FITC at pH 7.45. The number of GUVs used for the statistics is denoted by #GUVs and displayed in the 2nd column. The numerical values for the mean value and the standard deviation (SD) of the normalised intensity are dimentionless.

| $X$ [nM] | #GUVs | Mean coverage | SD |
|---|---|---|---|
| 0 | 3 | 0.014 | 0.004 |
| 15 | 4 | 0.347 | 0.088 |
| 30 | 4 | 0.585 | 0.050 |
| 60 | 4 | 0.707 | 0.184 |
| 120 | 4 | 0.748 | 0.085 |
| 180 | 4 | 0.822 | 0.127 |
| 240 | 4 | 0.818 | 0.075 |
| 480 | 4 | 0.834 | 0.027 |

Table S4: Statistics for the surface coverage of GUV membranes by 6H-FITC for three different preparation methods, corresponding to the data in Figure 7b. The membranes contained 3 mol% NTA lipids and were exposed to a 240 nM solution of 6H-FITC at pH 7.45. The number of GUVs used for the statistics is denoted by #GUVs and displayed in the 2nd column. The numerical values for the mean value and the standard deviation (SD) of the coverage are given in units of membrane-bound molecules per $\mu m^2$.

| Preparation method | #GUVs | Mean coverage | SD |
|---|---|---|---|
| Pt wire electroformation | 29 | 55 718 | 8 751 |
| PVA hydrogel swelling | 20 | 12 411 | 4 376 |
| ITO glass electroformation | 29 | 13 904 | 8 230 |

Table S5: Statistics for the fluorescence intensity of GUV membranes arising from membrane-bound 6H-GFP for different pH values in the exterior solution, corresponding to the data in Figure 9a. The membranes were prepared by platinum wire electroformation with 3 mol% NTA lipids and were exposed to a 20 nM solution of 6H-GFP. The pH was varied by the addition of HCl or NaOH solution. The number of GUVs used for the statistics is denoted by #GUVs. The numerical values for the mean value and the standard deviation (SD) of the coverage are given in units of membrane-bound molecules per $\mu m^2$.

| pH | #GUVs | Intensity | SD |
|---|---|---|---|
| 6.38 | 5 | 14.8 | 5.9 |
| 6.98 | 5 | 36.8 | 18.0 |
| 7.25 | 5 | 70.1 | 16.9 |
| 7.43 | 5 | 72.4 | 21.5 |
| 8.02 | 5 | 80.2 | 24.8 |
| 8.54 | 5 | 110.6 | 27.0 |
| 9.06 | 5 | 108.3 | 29.7 |
| 9.36 | 5 | 114.3 | 29.2 |

Table S6: Statistics for the fluorescence intensity of GUV membranes arising from membrane-bound 6H-FITC for different pH values in the exterior solution, corresponding to the data in Figure 9b. The membranes were prepared by platinum wire electroformation with 3 mol% NTA lipids and were exposed to a 120 nM solution of 6H-FITC. The pH was decreased and increased by the addition of HCl or NaOH solution. The number of GUVs used for the statistics is denoted by #GUVs. The numerical values for the mean value and the standard deviation (SD) of the coverage are given in units of membrane-bound molecules per $\mu m^2$.

| pH | #GUVs | Intensity | SD |
|---|---|---|---|
| 6.38 | 5 | 10.3 | 2.0 |
| 6.98 | 5 | 14.0 | 4.0 |
| 7.25 | 5 | 21.4 | 2.9 |
| 7.43 | 5 | 33.7 | 4.1 |
| 8.02 | 5 | 56.8 | 6.7 |
| 8.54 | 5 | 76.9 | 7.3 |
| 9.06 | 5 | 67.4 | 7.5 |
| 9.36 | 5 | 20.7 | 3.0 |

Table S7: Statistics for the fluorescence intensity of GUV membranes exposed to 6H-FITC for different electroformation methods and two different mole fractions of NTA lipids, corresponding to the data in Figure S2. The membranes were prepared with 3 mol% or 30 mol% NTA lipids and exposed to a 120 nM solution of 6H-FITC at pH 7.45. The number of GUVs used for the statistics is denoted by #GUVs. The numerical values for the mean value and the standard deviation of the coverage are given in units of membrane-bound molecules per $\mu m^2$.

| Preparation method | #GUVs | Intensity, 3 mol% | #GUVs | Intensity, 30 mol% |
|---|---|---|---|---|
| Pt wire electroformation | 17 | $34.4 \pm 4.9$ | 7 | $42.1 \pm 16.1$ |
| ITO glass electroformationn | 12 | $6.1 \pm 7.6$ | 6 | $17.8 \pm 7.3$ |

### Movie Caption

**Movie 1:** Time-lapse movie showing the exchange of 2 $\mu$M carboxyfluorescein solution to the side channels. The images are obtained by an overlay of the brightfield and the fluorescent channels. Initially, the device is filled with water and then the solution in the reservoir is exchanged with carboxyfluorescein. The pump connected to the outlet is drawing the solution at the speed of 150 $\mu$L/hr. The device is imaged close to the outlet. It takes around 3.5 mins to observe homogeneous fluorescence in the two main channels as well as in all side channels.
